# MALDI-IM-MS Imaging of Brain Sterols and Lipids in a Mouse Model of Smith-Lemli-Opitz Syndrome

**DOI:** 10.1101/2023.10.02.560415

**Authors:** Amy Li, Libin Xu

## Abstract

Smith-Lemli-Opitz syndrome (SLOS) is a neurodevelopmental disorder caused by genetic mutations in the *DHCR7* gene, which encodes the enzyme 3β-hydroxysterol-Δ^7^-reductase (DHCR7) that catalyzes the last step of cholesterol synthesis, resulting in deficiency in cholesterol and accumulation of its precursor, 7-dehydrocholesterol (7-DHC). To understand how the brain regions are differentially affected by the defective Dhcr7, we aim to map the regional distribution of sterols and other lipids in neonatal brains from a *Dhcr7*-KO mouse model of SLOS, using mass spectrometry imaging (MSI). MSI enables spatial localization of biomolecules *in situ* on the surface of a tissue section, which is particularly useful for mapping the changes that occur within a metabolic disorder such as SLOS, and in an anatomically complex organ such as the brain. In this work, using MALDI-ion mobility (IM)-MSI, we successfully determined the regional distribution of features that correspond to cholesterol, 7-DHC/desmosterol, and the precursor of desmosterol, 7-dehydrodesmosterol, in WT and *Dhcr7*-KO mice. Interestingly, we also observed *m/z* values that match the major oxysterol metabolites of 7-DHC (DHCEO and hydroxy-7-DHC), which displayed similar patterns to 7-DHC. We then identified brain lipids using *m/z* and CCS at the Lipid Species-level and curated a collection of MALDI-IM-MS-derived lipid CCS values. Subsequent statistical analysis of regions-of-interest allowed us to identify differentially expressed lipids between *Dhcr7*- KO and WT brains, which could contribute to defects in myelination, neurogenesis, neuroinflammation, and learning and memory in SLOS.

## 1. Introduction

Smith-Lemli-Opitz syndrome (SLOS) is a multiple malformation disorder caused by genetic mutations in the *DHCR7* gene, encoding the enzyme 3β-hydroxysterol-Δ^7^- reductase (DHCR7) that catalyzes the final step in cholesterol synthesis ^1, 2^. The biochemical hallmark of SLOS is cholesterol deficiency and accumulation of the precursor to cholesterol, 7-dehydrocholesterol (7-DHC), and during brain development, 7-dehydrodesmosterol (7-DHD) ^3-6^. Due to its high reactivity towards free radical oxidation ^7^, 7-DHC also gives rise to many other oxidized metabolites, known as oxysterols ^8-11^, among which 3β,5α-dihydroxycholest-7-en-6-one (DHCEO), 4α-hydroxy- 7-DHC, and 4β-hydroxy-7-DHC are the major ones observed in embryonic and postnatal day 0 (P0) *Dhcr7*-KO mouse brains ^6^. Oxysterols, whether produced enzymatically or non-enzymatically, are known to play both a pathological and physiological role in the brain and other tissues ^12-15^. Thus, the effects of blocking cholesterol synthesis at the level of DHCR7 can be viewed as two-fold: depletion of a critical cellular molecule, cholesterol, and accumulation of 7-DHC and its derived oxysterols to potentially toxic levels.

Individuals diagnosed with SLOS can exhibit a wide range of phenotypes across several organ systems due to the essential role of cholesterol during development ^16, 17^. The central nervous system (CNS) is one of the most severely affected organs in SLOS because the brain synthesizes cholesterol locally and independently. The CNS phenotype manifests as cognitive and behavioral deficits (ranging from autistic behaviors to severe developmental delay), as well as gross anatomical defects such as agenesis of the corpus callosum ^1^. This highlights the essential role of cholesterol in brain development. *De novo* synthesis of cholesterol begins early on in mouse embryonic development ^18^, and accumulation of 7-DHC-derived oxysterols has been observed as early as E12.5 in *Dhcr7*-KO mouse brain ^6^. This early accumulation of 7-DHC-derived oxysterols in SLOS was shown to be toxic to neuronal cells and cause decreased proliferation and premature differentiation of neural precursor cells ^6, 19^.

The *Dhcr7*^*tm1Gst*^ (*Dhcr7*^*Ex8*^) mouse model of SLOS is equivalent to the IVS8-1G>C in humans and is associated with a severe phenotype, causing postnatal lethality within 24 hours of birth in homozygous mice. Previously, we have conducted an untargeted lipidomics study of the brain over the time course of development in this mouse model of SLOS using hydrophilic interaction liquid chromatography-ion mobility-mass spectrometry (HILIC-IM-MS) ^20^. While the study was informative on global changes in various lipid classes and species as embryonic development progressed, whole brain tissue homogenization eliminated the rich spatial information present within the brain. Matrix-assisted laser desorption ionization (MALDI)-based mass spectrometry imaging (MSI) allows spatial information of endogenous metabolites to be preserved and is thus well suited for studying the changes in the distribution and abundance of various brain lipids in the SLOS mouse model. This study represents a crucial step in understanding how the brain regions are differentially affected by the loss-of-function mutation in *Dhcr7* and the role of lipid metabolism in the overall pathophysiology of SLOS.

However, relative to LC-based lipidomics, MALDI lacks the chromatographic separation of analytes, which presents a challenge for the identification of lipids because most lipids are observed in a narrow mass range of 600-900 Dalton. Coupling MALDI with IM-MS provides an additional dimension of separation of lipids without significantly decreasing the throughput. IM separates ions based on their collision cross sections (CCS) with an inert background gas, which is dependent on the size and shape of the ions and the specific gas. It has been demonstrated that different classes of lipids tend to occupy different chemical spaces in the CCS-*m/z* plot ^21-25^, which can be used to enhance the confidence of lipid identification.

In this work, we aim to map the regional distribution of sterols and lipids using MALDI-IM-MS. We first measured the CCS values of ions derived from sterols and oxysterols and successfully determined the distribution of cholesterol and its precursors as well as the *m/z* values that correspond to the oxysterol metabolites of 7-DHC. We then identified lipids using *m/z* and CCS at the Lipid Species-level (lipid class/subclass and fatty acid sum composition) according to the Lipidomics Standards Initiative ^26^. Subsequent statistical analysis revealed differentially expressed lipids between *Dhcr7*- KO and WT mouse brains.

## 2. Methods and Materials

### 2.1 Chemicals

α-Cyano-hydroxycinnamic acid (CHCA), 9-aminoacridine (9-AA), red phosphorus, and cholesterol were purchased from Sigma-Aldrich Inc. 2-Methylbutane was purchased from Thermo Fisher Scientific (Grand Island, New York). 7-dehydrocholesterol, 7- dehydrodesmosterol, and desmosterol were purchased from Avanti Polar Lipids. DHCEO, 4α-hydroxy-7-DHC, and 4β-hydroxy-7-DHC were prepared as described previously ^9, 10^.

### 2.2 Animals

C57BL/6J and transgenic heterozygous mice with a null mutation for *Dhcr7* (Ex8) mice were purchased from Jackson Laboratories (Bar Harbor, Maine; catalogue #007453). Studies reported in this work were approved by the University of Washington Institutional Animal Care and Use Committee. Mice were housed in an animal care facility with a 12-hour light and dark cycle and fed an *ad libitum* commercial rodent chow diet. Heterozygous *Dhcr7* (Ex8) mice were mated overnight, where the day after time-mating was designated as embryonic day 0.5 (E0.5). At birth, neonate litters were euthanized and their brains were harvested under a dissection scope and flash frozen in pre-cooled 2-methylbutane, transferred to dry ice, and stored at -80ºC until further steps of sectioning, matrix application, and analysis.

### 2.3 Sample preparation for MALDI-MSI

The sample preparation workflow is outlined in **Figure 1**. Brain tissues were attached to the cryostat chuck with minimal OCT compound. Coronal tissue sections were cut at 16 µm thickness and thaw-mounted onto standard glass slides. Sections from WT and KO mouse brains were collected at various intervals along the anterior-posterior axis and paired together on the same slide. Slides were stored at -80 ºC and placed in a vacuum desiccator to thaw to avoid condensation.

**Figure 1.**
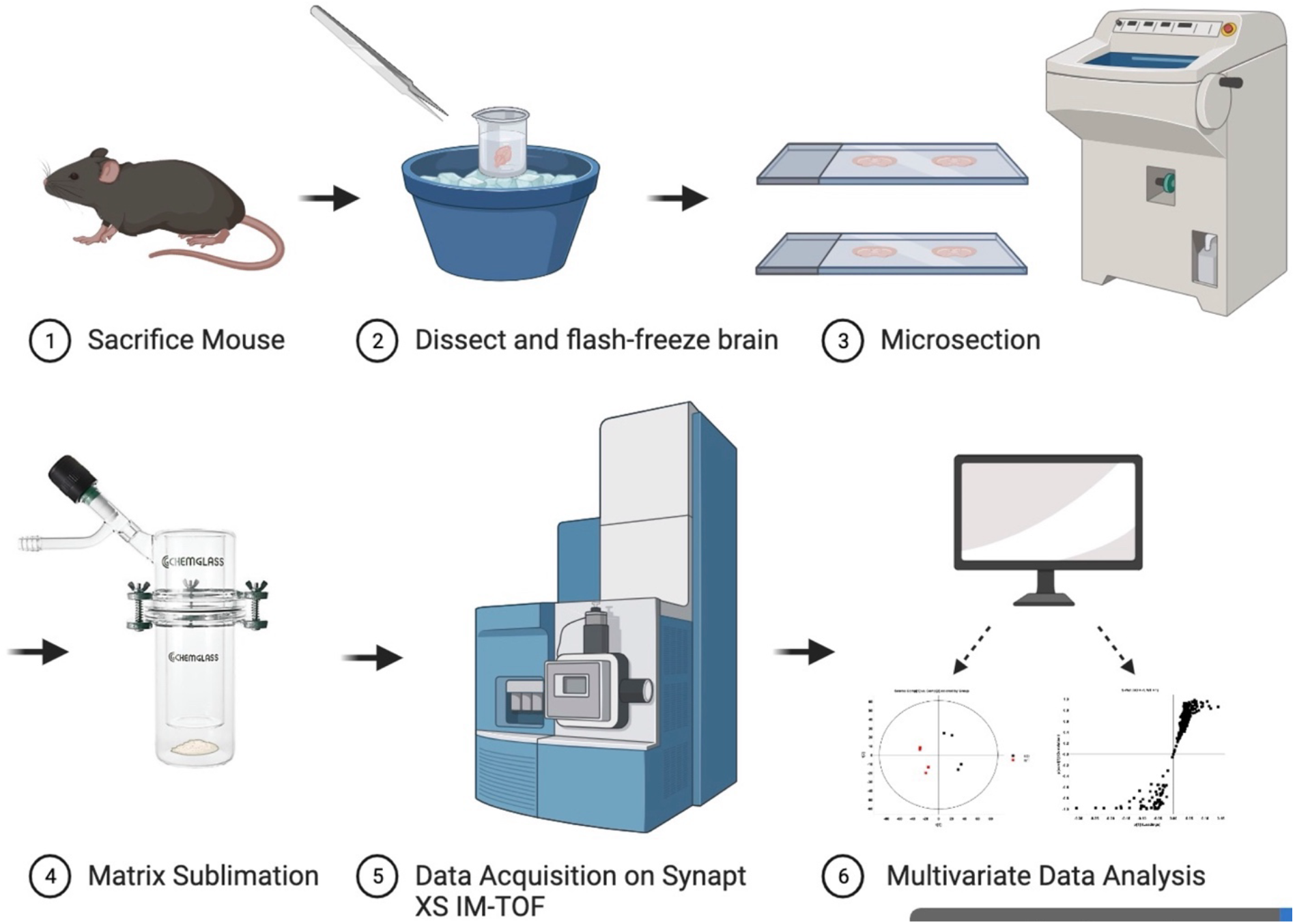
Diagram of the MALDI-IM-MS imaging workflow, including steps for sample preparation (brain harvested from mouse, tissue sectioning), matrix application (coating via sublimation), and data acquisition and analysis.

Slides were coated *via* sublimation with CHCA and 9-AA for positive and negative mode ionization, respectively ^27^. Briefly, 300 mg of the matrix was evenly deposited on the bottom of the outer flask of the sublimation glassware (Chemglass Life Sciences), and a slide was affixed with tape to the bottom of the inner flask filled with an ice water slurry. The assembled glassware apparatus was heated up in a sand bath and connected to a vacuum pump manifold. Sublimation was performed under the following conditions: 180ºC, 0.2 Torr pressure, and 60 minutes for CHCA, and 120ºC, 2 Torr pressure, and 10 minutes for 9-AA.

H&E staining was performed on adjacent serial sections to the imaged sections following a standard H&E procedure (Vector Laboratories) to correlate ion images with specific histological brain regions. High-resolution optical images were obtained with a scanner (Epson Perfection V850 Pro).

### 2.4 MALDI-IM-MS

MALDI-IM-MS was performed on a Waters SYNAPT XS traveling wave IM (TWIM)-QTOF mass spectrometer equipped with a 1 kHz Nd:YAG laser in both positive and negative ionization modes. Data was acquired over a mass range of 50 to 1200 *m/z*. TWIM separation was performed with a gas flow rate of 90 mL/min, a wave velocity of 500 m/s, and a wave height of 40 V. Images were acquired at 30-50 µm spatial resolution. High-Definition Imaging (HDI) software and EZ-Info (Waters) were used for data processing, visualization, and statistical analysis. TWIM-derived CCS values were calibrated manually with a series of phosphatidylcholine (PC) and phosphatidylethanolamine (PE) lipid standards using an in-house Python script as described previously ^28, 29^. Red phosphorus reference spots were used for lock-mass correction. Features were normalized to the total ion current (TIC) at each pixel. Lipid identifications were made by *m/z* and CCS matching using various databases, including LIPID MAPS, HMDB, our own in-house CCS database, CCSbase (https://CCSbase.net)^30^, and an automated lipidomics data processing Python package, *LiPydomics* (https://github.com/dylanhross/lipydomics) ^22^. Lipid identifications are reported at the level of fatty acyl sum composition in **Supplementary Table S1**. On-tissue targeted MS/MS fragmentation of high-abundance lipids was carried out by ramping the collision energy in the transfer region from 25-40 eV.

Data shown in this work are from two biological replicates per ionization mode (replicate data shown in **Supplementary Figures S2-5)**.

## 3. Results

### 3.1 Confirmation of biochemical hallmark in *Dhcr7*-KO mouse brains

The biochemical hallmarks of *Dhcr7*-KO in the SLOS mouse brains, compared to the wild-type, are decreased levels of cholesterol and increased levels of 7-DHC and 7-DHD ^6^. To facilitate the identification of these sterols, we first analyzed standards of cholesterol, three precursor sterols (desmosterol, 7-DHC, and 7-DHD), and three major 7-DHC-derived oxysterols (DHCEO, 4α-hydroxy-7-DHC, and 4β-hydroxy-7-DHC), previously observed in *Dhcr7*-KO mouse brains,^6^ using MALDI-IM-MS in the positive mode with CHCA as the matrix. The *m/z* and CCS values of major peaks found from authentic standards are shown in **Figure 2A**, which are mostly [M+H-H_2_O]^+^ or further dehydration ions, as well as dehydrogenation ions for 7-DHC and 7-DHD. For sterols, major peaks correspond to the loss of one or two water molecules from the molecular ion ([M+H-H_2_O]^+^ or [M+H-2H_2_O]^+^), and for 7-DHC and 7-DHD, a dehydrogenation ion ([M+H-H_2_O-2H]^+^) at *m/z* 365.3 and 363.3, respectively, was also observed. We note that *m/z* 367.3 is the primary ion formed from both 7-DHC and desmosterol, which complicates the analysis. However, the *m/z* 365.3 is specific to the dehydrogenation ion of 7-DHC, [M+H-H_2_O-2H]^+^, and the dehydration ion of 7-DHD, [M+H-H_2_O]^+^. Although these two isomeric ions were not resolved by TWIM, they can still be used to inform the characteristic sterol distribution in the SLOS brain.

**Figure 2.**
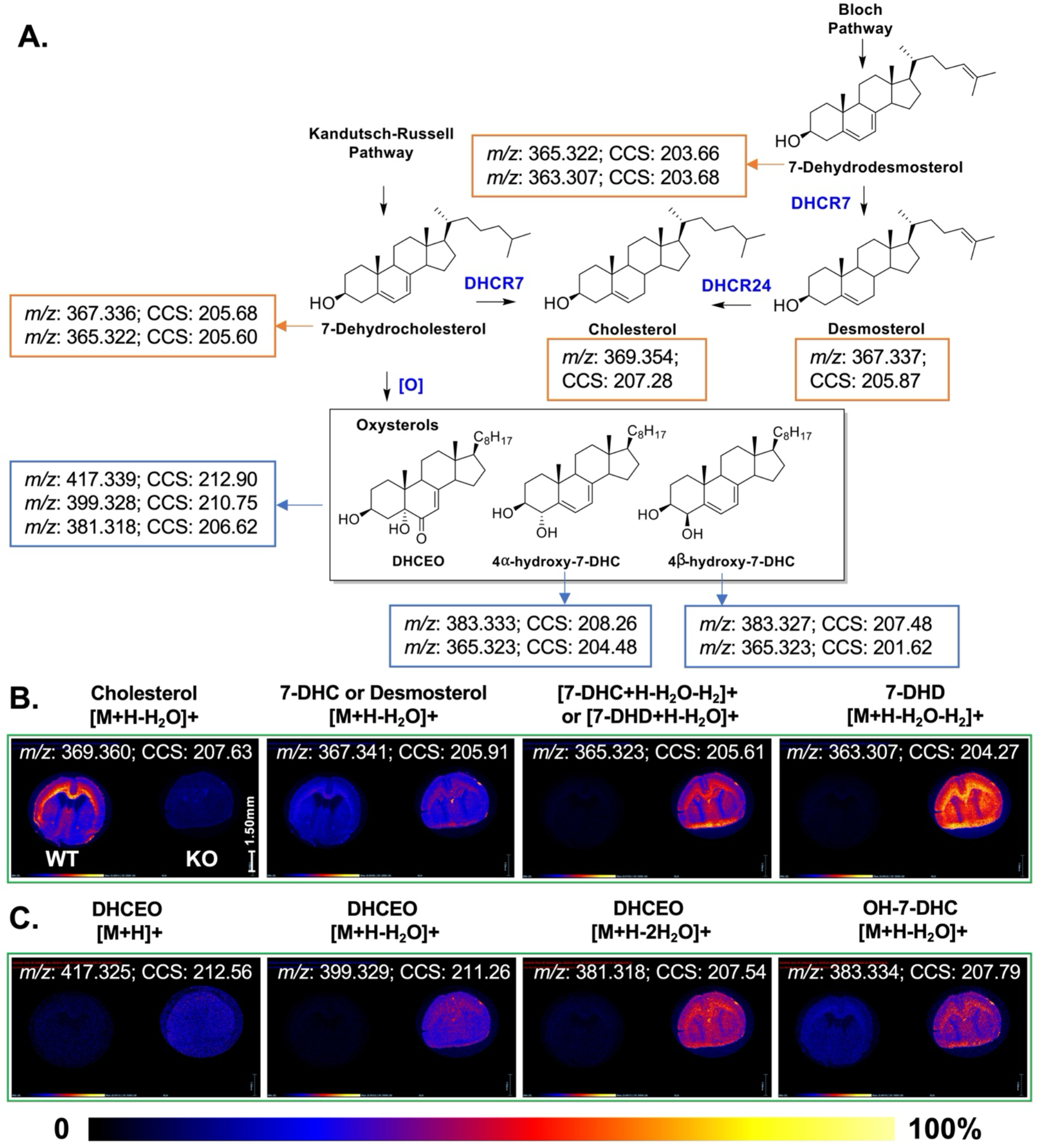
(A) Last steps of cholesterol biosynthesis pathway with associated enzymes. Boxes contain exact mass and TWIM-derived CCS measurements from MALDI of sterol and oxysterol standards. (B) Ion images of cholesterol and sterol precursors in WT (left) and *Dhcr7*-KO (right) mouse brain tissue. (C) Ion images of oxysterol metabolites in WT (left) and *Dhcr7*-KO (right) mouse brain tissues. Intensities are normalized to TIC and represented according to the heat map color gradient.

For oxysterols, the molecular ion of DHCEO ([M+H]^+^) was observed at *m/z* 417.3, as well as peaks for the loss of one or two water molecules (*m/z* 399.3 and 381.3). For 4α-hydroxy-7-DHC and 4β-hydroxy-7-DHC, only the loss-of-water ions were observed at *m/z* 383.3 and 365.3. These fragmentation patterns are consistent with those observed with atmospheric pressure chemical ionization on these oxysterols ^9, 10^.

Using MALDI-IM-MSI, we were able to investigate patterns of distribution of several endogenous sterols and oxysterols in the brain tissue of WT and *Dhcr7*-KO mice, as shown via ion images (**Figure 2B**). The heat map gradient shows the signal intensity of individual pixels within the tissue map. Anatomically, cholesterol is abundant in the corpus callosum structure of the WT mouse, which would be expected for such brain structures that are highly abundant in white matter. On the other hand, the signal for cholesterol is at a very low intensity on the *Dhcr7*-KO mouse tissue section. The ion image for *m/z* 367, which could arise from 7-DHC in the *Dhcr7*-KO brain and desmosterol in the WT brain, is more intense in the KO than the WT, but not completely absent in the WT due to the presence of desmosterol. On the other hand, the ions at *m/z* 365 and 363 are highly abundant in *Dhcr7*-KO tissue while virtually absent in the WT brain. The ion image for *m/z* 365 is most likely the combined signal from [M+H-H_2_O]^+^ of 7-DHD or the dehydrogenation ion of 7-DHC, [M+H-H_2_O-2H]^+^, while *m/z* 363 could result from the dehydrogenation ion of 7-DHD. The localization pattern of cholesterol precursors in the *Dhcr7*-KO tissue appears to be more diffuse compared with that of the cholesterol in the WT tissues. Although the signals are still the most intense in the corpus callosum, they are also abundant in the outer layer of the cortex, ventricular/subventricular zone, and brainstem.

Patterns for ion images of *m/z* corresponding to oxysterol compounds (**Figure 2C**), including three peaks (*m/z* 417, 399, and 381) that correspond to DHCEO ions, are similar to those of 7-DHC and 7-DHD. The molecular ion at *m/z* 417 shows the least amount of signal, while the loss-of-water ions, *m/z* 399 and 381, display good intensities. *m/z* 383 could arise from hydroxylated 7-DHC, which matches the exact mass and CCS values of our pure standards of 4α-hydroxy-7-DHC and 4β-hydroxy-7-DHC. However, their low abundances precluded any further investigation via fragmentation to obtain information on specific isomers.

### 3.2 Lipid CCS observed in mouse brains with MALDI

As expected, sterol and oxysterol features showed drastic changes in *Dhcr7*-KO brains compared with WT (**Figure 2**). However, we are also interested in other lipid species changes due to the lipidomic changes that can result from KO of *Dhcr7*.^20^ To accomplish this, we first aimed to identify brain lipids observed under MALDI using a combination of *m/z* and CCS. TWIM-derived CCS values of observed features were calibrated with a series of PC and PE standards for positive and negative modes, respectively, as described previously ^28^. Lipids were first identified using *LiPydomics* by setting the tolerance for *m/z* to 30 ppm and CCS to 3%. Unidentified features were further checked against *m/z* and CCS values in HMDB, which originate from AllCCS ^31^. Specifically, sterols, PCs, lyso-PCs (LPCs), ceramides (Cer), and diglycerides (DGs) were identified in the positive mode, while fatty acids (FAs), phosphatidylethanolamines (PEs), phosphatidylserines (PSs), phosphatidylinositols (PIs), phosphatidylinositol phosphates (PIPs), phosphatidylglycerols (PGs), phosphatidic acids (PAs), lyso-PAs (LPAs), lyso-PEs (LPEs), lyso-PGs (LPGs), lyso-PSs (LPSs), and lyso-PIs (LPIs) were identified in the negative mode. **Supplementary Table 1** and **Figure 3** summarize all the identified lipids at the lipid species level with the sum fatty acid composition. As seen in **Figure 3**, lipid distribution in the CCS-*m/z* plot is similar to what is observed with ESI ^21^. In the positive mode CCS-*m/z* plot (**Figure 3A**), the CCS trend of sterol metabolites is lower than that of phospholipids and sphingolipids, suggesting that they adopt more compact gas-phase conformations due to their fused-ring structures.

**Figure 3.**
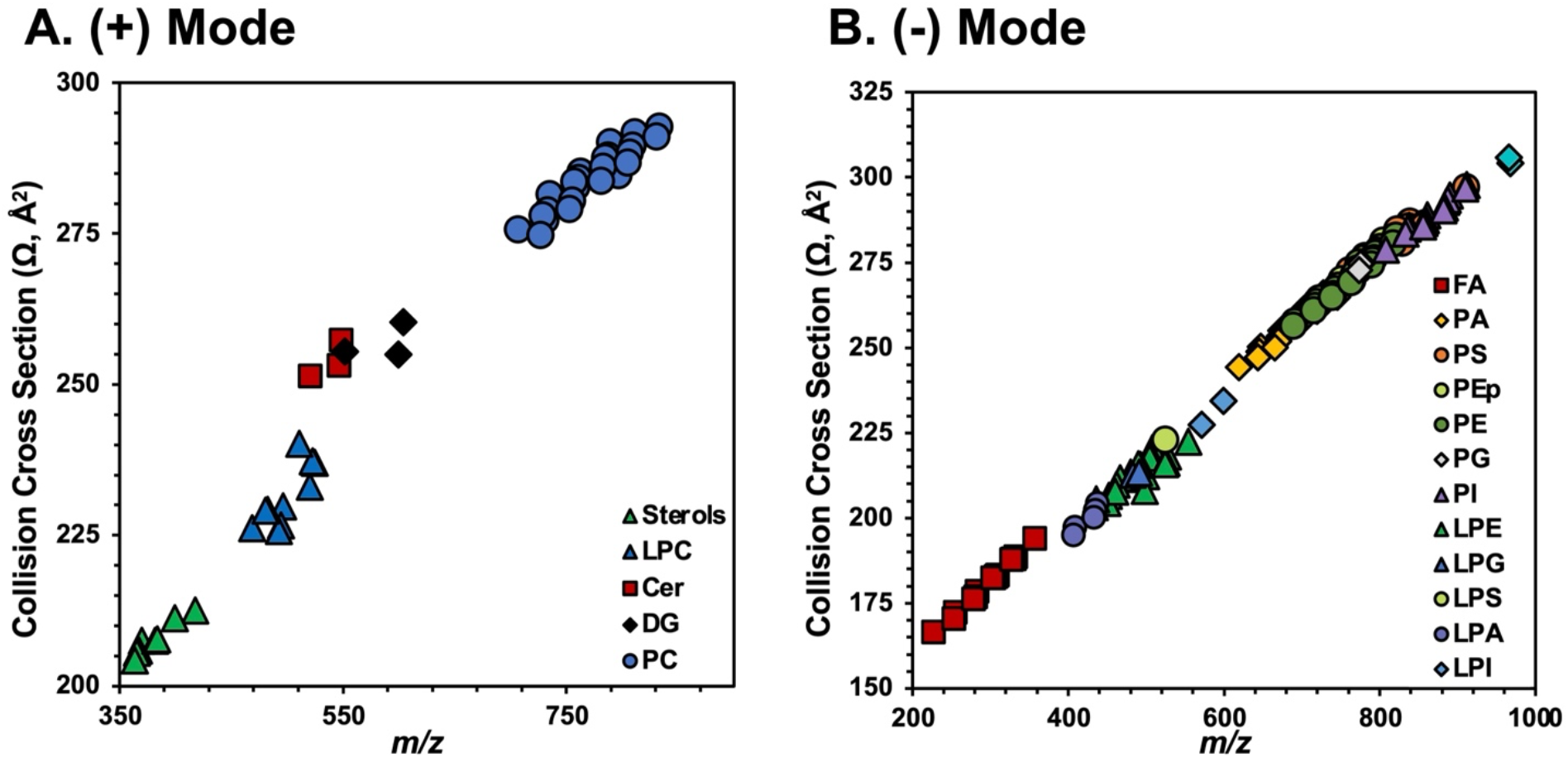
Collision cross section (CCS)-*m/z* plot of lipids identified in (A) positive and (B) negative ionization modes in MALDI-IM-MS analysis of mouse brains at postnatal day 0.

From these data, some lipid species with isobaric *m/z* values but large differences in CCS can be found. For example, *m/z* values of [M-H]^-^ ions of LPE(22:6) and LPS(18:0) differ by only 0.004%, but their CCS values differ by 3.3%. As another example, the *m/z* values of [M+Cl]^-^ ion of PC(34:1) and the [M-H]^-^ ion of PE(40:4) differ by 0.002%, but their CCS values differ by 3.3%. Other isobaric pairs of lipids with different CCS are listed in **Supplementary Table S2**. Images of selected isobaric pairs are shown in **Figures 4** and **S3**, which demonstrate distinct regional distribution between these pairs. Thus, CCS enabled more confident identification and imaging of isobaric lipids.

**Figure 4.**
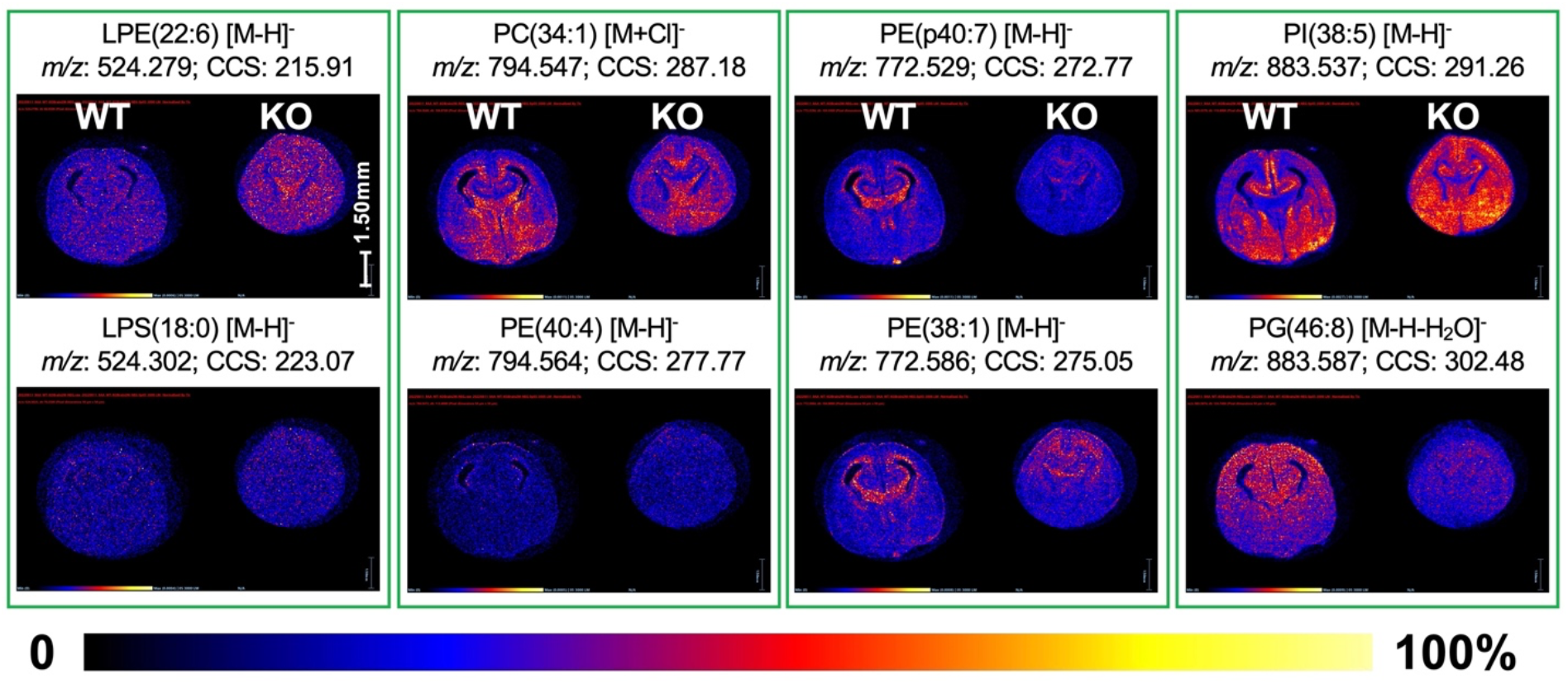
Ion images of selected isobaric lipid pairs with different CCS values. Intensities are normalized to TIC and represented according to the heat map color gradient.

### 3.3 Differentially expressed lipids between WT and *Dhcr7*-KO mouse brains

Using multivariate statistical analysis tools, we identified lipid species that were differentially expressed in the *Dhcr7*-KO mouse brains compared to WT. Principal component analysis (PCA) plots for regions-of-interest (ROI) from WT and *Dhcr7*-KO brain sections (four quadrants from one tissue section) show clustering of and separation between the two genotypes along PC1 (**Supplementary Figure S1**). S-plots generated from OPLS-DA (**Figure 5A**) revealed differential lipid features that drive the separation between WT and *Dhcr7-*KO brains, where p[1] on the x-axis, describes how much a feature influences the separation on the OPLS-DA, while p(corr) on the y-axis, measures the reliability of that feature as a contributing factor in the model. Features with p(corr) greater than 0.9 or less than -0.9 are denoted with an asterisk in **Supplementary Table 1**. Selective features that display a high correlation with WT (lower left) or KO (upper right) are displayed in **Figure 5B** along with the H&E staining of adjacent serial sections. Some features were present only in either WT or *Dhcr7-*KO tissue, and others that showed more subtle changes were picked up via S-plot analysis. Additional replicates of the images from a different brain are shown in **Figure S4**.

**Figure 5.**
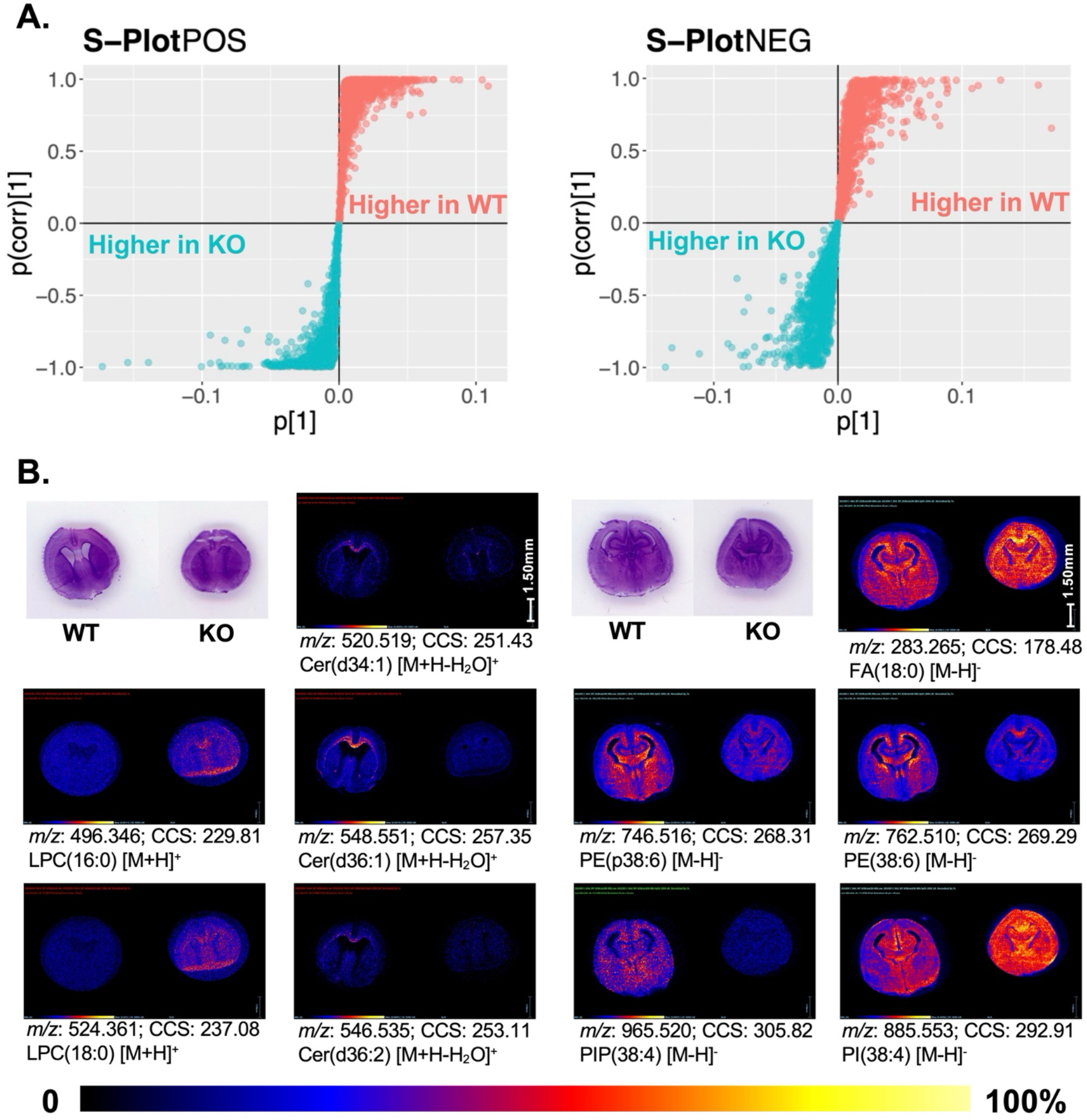
(A) S-plots from multivariate statistical analysis of a pair of WT and *Dhcr7*-KO mouse brain tissues in positive and negative ionization modes. (B) Ion images of lipid features that are higher in WT or higher in *Dhcr7*-KO are shown below each respective S-plot. Intensities are normalized to TIC and represented according to the heat map color gradient.

**Figure 6.**
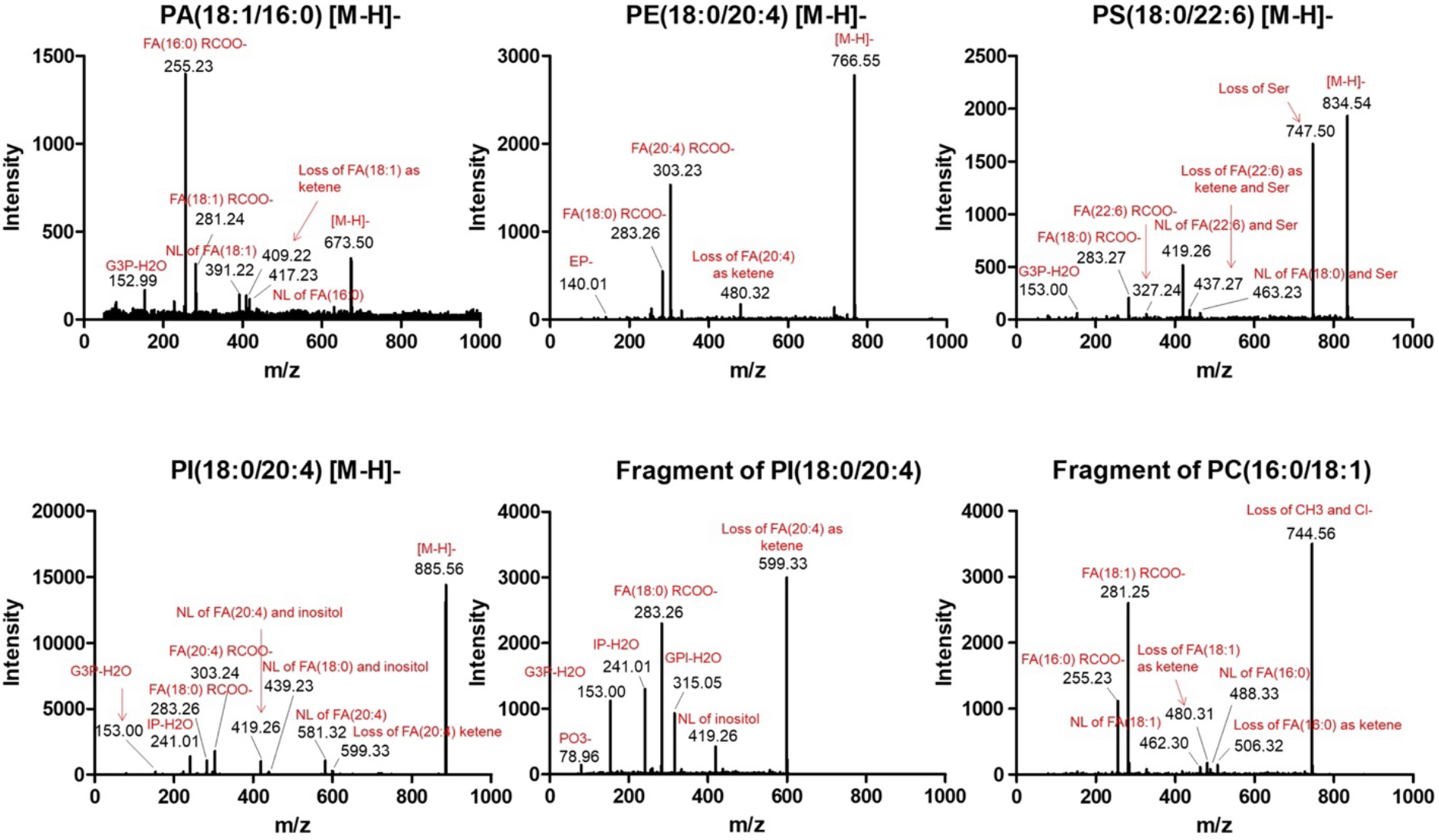
Targeted fragmentation spectra of lipid features found on tissue via MALDI-MS/MS. Diagnostic fragments were used to confirm lipid class assignment and determine sn1/sn2 fatty acyl chain compositions.

Our previous HILIC-IM-MS study highlighted certain changes in lipids across developmental timepoints ^20^, and we were interested to see if similar changes would be detected via MALDI-IM-MSI, but with a spatial resolution component. ESI and MALDI are two fundamentally different ionization sources, which could lead to different observations of lipids and differences due to ion suppression of low-abundant lipids when analyzing the whole brain homogenates. We observed significantly decreased levels in several ceramides and significantly increased levels in lysoPC and lysoPE species in the KO samples relative to WT (**Figure 5** and **Table S1**), consistent with the lipidomic changes identified using HILIC-IM-MS. The ceramides were most abundant in the corpus callosum of WT (made up of white matter) and appeared to be mostly absent in *Dhcr7*-KO. Another interesting observation was PI species detected in the negative ionization mode, where PI(38:4) is upregulated in *Dhcr7*-KO, while the corresponding PI monophosphate, PIP(38:4), is downregulated in *Dhcr7*-KO tissue, which could point to aberrant PI signaling pathways. In general, we also observed increased levels of free fatty acids (FFA) in the *Dhcr7*-KO brains, such as FFA 18:1, 20:3, 20:4, and 22:3. Images of additional lipid species are shown in **Supplementary Figure S5**.

Ions with similar tissue distributions could be identified using the image correlation feature in HDI. The ion at *m/z* 660 (**Supplementary Figure S5**) had a very high correlation with the cholesterol peak, *m/z* 369. This and other unknown features that had a stark contrast between *Dhcr7*-KO and WT tissues were not identified by any databases. Low abundance of these lipids precluded targeted fragmentation to obtain more structural information.

### 3.4 On-tissue MALDI-MS/MS confirms abundant lipid annotations

Some of the more abundant lipid species were confirmed via on-tissue targeted MS/MS fragmentation in the negative ionization mode. Fragmentation spectra were assigned in **Figure 5** using the LIPID MAPS prediction tool for the fragmentation of phospholipids. Diagnostic fragments were used to confirm the lipid class and determine the specific sn1/sn2 position of fatty acids. Thus, targeted fragmentation spectra of PE(18:0/20:4), PS(18:0/22:6), and PI(18:0/20:4) showed diagnostic fragments of ethanolamine phosphate at *m/z* 140.01, loss of serine headgroup at *m/z* 747.50, inositol phosphate dehydration ion at *m/z* 241.01, respectively.

Additionally, fragmentation revealed that two specific *m/z* features were in-source fragments of other abundant lipids, including the fragment of PI(18:0/20:4), which had the characteristic peak of inositol phosphate, and the fragment of PC(16:0/18:1), which had diagnostic fragments that included the intact choline headgroup.

Additionally, PA(18:1/16:0) could also be an in-source fragment, and this has been reported in the literature as a potential source of error in MALDI experiments ^32^. Any headgroup loss due to in-source fragmentation would lead to corresponding spectra for PA and thus, could be mistakenly identified as a PA because there are no unique diagnostic fragments for PA.

## 4. Discussion

In this work, we successfully established the spatial distribution of cholesterol and its precursors, 7-DHC and 7-DHD, in the WT and *Dhcr7*-KO mouse model of SLOS. It has been previously reported that 7-DHC-derived oxysterols, particularly DHCEO, cause the neurogenic defects observed in SLOS models ^6, 19^. While increased levels of these oxysterols have been reported previously in whole-brain homogenates, this work represents the first molecular imaging of oxysterols in developing mouse brains. We found that the oxysterols mostly co-localized with the images of *m/z* values that correspond to 7-DHC and 7-DHD, with the highest intensity observed in the corpus callosum, followed by the outer layer of the cortex, ventricular/subventricular zone, and brainstem. Because neurogenesis initiates in the ventricular regions, the images support the roles of the cholesterol precursors and the oxysterols in mediating the defective neurogenesis in the *Dhcr7*-KO mouse model that was previously reported.^6^ The high level of sterols in the corpus callosum is also consistent with the commonly observed defects in the corpus callosum in the brains of SLOS patients.^1, 33^

This work also assembled a collection of lipid CCS values (48 and 134 values in the positive and negative modes, respectively) derived from MALDI and TWIM. Thus, this lipid CCS collection fills in a gap in MALDI-derived CCS values since previous CCS values are mostly measured for ions formed from ESI. Significantly, although MALDI and ESI are fundamentally different ionization mechanisms, the lipid CCS values are consistent between the same ions formed from MALDI and ESI. This demonstrates that the widely available ESI-derived CCS library ^23, 30, 31, 34^ can be used for metabolite identification when using MALDI.

Differentially expressed lipids in different brain regions between *Dhcr7*-KO and WT mouse brains could shed light on their contribution to SLOS pathophysiology, in addition to sterols and oxysterols. For example, the decreased levels of ceramides in the corpus callosum could affect the myelination process because ceramides are the lipid precursor to sphingomyelin, which is a major lipid involved in myelination. White matter lesions have been observed in the brains of SLOS patients,^33, 35^ which could be attributed to myelination defects. Second, the increased level of PI(38:4) and decreased level of PIP(38:4) could have a profound impact on the neurogenesis process.

Conversion of PI to PIP is mediated by phosphatidylinositol 3-kinase (PI3K), and the PI3K/AKT pathway has been found to be important for neurogenesis ^36^. Third, lysophospholipids were found to be increased in *Dhcr7*-KO brains. Lysophospholipids are important signaling molecules that mediate numerous physiological processes, such as brain growth, neuron arborization, and myelination, through their specific G-protein coupled receptors ^37^. Lastly, free fatty acids were, in general, higher in the *Dhcr7*-KO brains than in the WT, which is consistent with the higher level of lysophospholipids since both are presumably formed from the phospholipase activities. Free fatty acids have been found to play important roles in neuroinflammation, learning, and memory.^38^ To summarize, the spatial lipidomic changes resulting from *Dhcr7*-KO could lead to new avenues of research to further understand the neurodevelopmental phenotype in SLOS.

## Conclusion

In this work, we used MALDI-IM-MSI to map out the brain regional sterolomic and lipidomic changes caused by *Dhcr7*-KO in a SLOS mouse model compared to wild-type controls. Although cholesterol is a notoriously difficult molecule to ionize, we were able to achieve mapping of the distribution of cholesterol, sterol precursors, and oxysterol metabolites within WT and *Dhcr7*-KO brains in positive ionization mode, which confirmed the biochemical hallmark of SLOS. In addition, we found that several classes of lipids in brain tissue were altered between WT and *Dhcr7*-KO via multivariate statistical analysis, and confirmed their identities via *m/z* and CCS. On-tissue fragmentation further confirmed the identities and fatty acid composition of some abundant lipids. The alterations in ceramides, lysophospholipids, and phosphatidylinositols could have important implications in myelination, neural differentiation, and neurogenesis in the brain with *Dhcr7* deficiency.

## Data availability

Raw MALDI-IM-MS data is available at MassIVE under MSV000092638 (doi:10.25345/C5125QM05).

## Acknowledgment

This work was supported by a National Institutes of Health (NIH) grant R01HD092659. AL was supported by the UW Pharmacological Sciences Training Program (NIH T32 GM007750) and the Institute of Translational Health Sciences TL1 Program (NIH TL1 TR002318).

## Conflict of Interest statement

The authors declare that they have no conflicts of interest with the contents of this article.

## Notes

### Competing Interest Statement

The authors have declared no competing interest.

### Summary of Updates

We updated Supplemental Tables S1 and S2 and Figures 2, 3, 5, and S4. Some related text was also updated.

## References

1. Porter, F. D.; Herman, G. E., Malformation syndromes caused by disorders of cholesterol synthesis. J. Lipid Res. 2011, 52 (1), 6–34.

2. Thurm, A., et al., Development, behavior, and biomarker characterization of Smith-Lemli-Opitz syndrome: an update. J. Neurodev. Disord. 2016, 8, 12.

3. Irons, M.; Elias, E. R.; Salen, G.; Tint, G. S.; Batta, A. K., Defective cholesterol biosynthesis in Smith-Lemli-Opitz syndrome. Lancet 1993, 341 (8857), 1414.

4. Tint, G. S.; Irons, M.; Elias, E. R.; Batta, A. K.; Frieden, R.; Chen, T. S.; Salen, G., Defective cholesterol biosynthesis associated with the Smith-Lemli-Opitz syndrome. N. Engl. J. Med. 1994, 330 (2), 107–113.

5. Tint, G. S.; Seller, M.; Hughes-Benzie, R.; Batta, A. K.; Shefer, S.; Genest, D.; Irons, M.; Elias, E.; Salen, G., Markedly increased tissue concentrations of 7-dehydrocholesterol combined with low levels of cholesterol are characteristic of the Smith-Lemli-Opitz syndrome. J. Lipid Res. 1995, 36 (1), 89–95.

6. Tomita, H.; Hines, K. M.; Herron, J. M.; Li, A.; Baggett, D. W.; Xu, L., 7-Dehydrocholesterol-derived oxysterols cause neurogenic defects in Smith-Lemli-Opitz syndrome. eLife 2022, 11, e67141.

7. Xu, L.; Davis, T. A.; Porter, N. A., Rate Constants for Peroxidation of Polyunsaturated Fatty Acids and Sterols in Solution and in Liposomes. J. Am. Chem. Soc. 2009, 131 (36), 13037–13044.

8. Xu, L.; Korade, Z.; Porter, N. A., Oxysterols from free radical chain oxidation of 7-dehydrocholesterol: product and mechanistic studies. J. Am. Chem. Soc. 2010, 132 (7), 2222–2232.

9. Xu, L.; Korade, Z.; Rosado, D. A.; Liu, W.; Lamberson, C. R.; Porter, N. A., An oxysterol biomarker for 7-dehydrocholesterol oxidation in cell/mouse models for Smith-Lemli-Opitz syndrome. J. Lipid Res. 2011, 52 (6), 1222–1233.

10. Xu, L.; Liu, W.; Sheflin, L. G.; Fliesler, S. J.; Porter, N. A., Novel oxysterols observed in tissues and fluids of AY9944-treated rats - a model for Smith-Lemli-Opitz Syndrome. J. Lipid Res. 2011, 52 (10), 1810–1820.

11. Xu, L.; Korade, Z.; Rosado, D. A., Jr.; Mirnics, K.; Porter, N. A., Metabolism of oxysterols derived from nonenzymatic oxidation of 7-dehydrocholesterol in cells. J. Lipid Res. 2013, 54 (4), 1135–43.

12. Bjorkhem, I.; Cedazo-Minguez, A.; Leoni, V.; Meaney, S., Oxysterols and neurodegenerative diseases. Mol Aspects Med 2009, 30 (3), 171–179.

13. Brown, A. J.; Jessup, W., Oxysterols and atherosclerosis. Atherosclerosis 1999, 142 (1), 1–28.

14. Raleigh, D. R., et al., Cilia-Associated Oxysterols Activate Smoothened. Mol Cell 2018, 72 (2), 316–327 e5.

15. Sacchetti, P., et al., Liver X receptors and oxysterols promote ventral midbrain neurogenesis in vivo and in human embryonic stem cells. Cell Stem Cell 2009, 5 (4), 409–19.

16. Porter, J. A.; Young, K. E.; Beachy, P. A., Cholesterol modification of hedgehog signaling proteins in animal development. Science 1996, 274 (5285), 255–9.

17. Cooper, M. K.; Wassif, C. A.; Krakowiak, P. A.; Taipale, J.; Gong, R.; Kelley, R. I.; Porter, F. D.; Beachy, P. A., A defective response to Hedgehog signaling in disorders of cholesterol biosynthesis. Nat. Genet. 2003, 33 (4), 508–13.

18. Tint, G. S.; Yu, H.; Shang, Q.; Xu, G.; Patel, S. B., The use of the Dhcr7 knockout mouse to accurately determine the origin of fetal sterols. J. Lipid Res. 2006, 47 (7), 1535–41.

19. Xu, L.; Mirnics, K.; Bowman, A. B.; Liu, W.; Da, J.; Porter, N. A.; Korade, Z., DHCEO accumulation is a critical mediator of pathophysiology in a Smith-Lemli-Opitz syndrome model. Neurobiol. Dis. 2012, 45 (3), 923–9.

20. Li, A.; Hines, K. M.; Ross, D. H.; MacDonald, J. W.; Xu, L., Temporal changes in the brain lipidome during neurodevelopment of Smith-Lemli-Opitz syndrome mice. Analyst 2022, 147 (8), 1611–1621.

21. Hines, K.; Herron, J.; Xu, L., Assessment of Altered Lipid Homeostasis by HILIC-Ion Mobility-Mass Spectrometry-Based Lipidomics. J. Lipid Res. 2017, 58 (4), 809–819.

22. Ross, D. H.; Cho, J. H.; Zhang, R.; Hines, K. M.; Xu, L., LiPydomics: A Python Package for Comprehensive Prediction of Lipid Collision Cross Sections and Retention Times and Analysis of Ion Mobility-Mass Spectrometry-Based Lipidomics Data. Anal. Chem. 2020, 92, 14975–.

23. Zhou, Z.; Tu, J.; Xiong, X.; Shen, X.; Zhu, Z. J., LipidCCS: Prediction of Collision Cross-Section Values for Lipids with High Precision To Support Ion Mobility-Mass Spectrometry-Based Lipidomics. Anal. Chem. 2017, 89 (17), 9559–9566.

24. Leaptrot, K. L.; May, J. C.; Dodds, J. N.; McLean, J. A., Ion mobility conformational lipid atlas for high confidence lipidomics. Nat Commun 2019, 10 (1), 985.

25. Kyle, J. E., et al., Uncovering biologically significant lipid isomers with liquid chromatography, ion mobility spectrometry and mass spectrometry. Analyst 2016, 141 (5), 1649–59.

26. Liebisch, G.; Vizcaino, J. A.; Kofeler, H.; Trotzmuller, M.; Griffiths, W. J.; Schmitz, G.; Spener, F.; Wakelam, M. J., Shorthand notation for lipid structures derived from mass spectrometry. J. Lipid Res. 2013, 54 (6), 1523–30.

27. Sarah Caughlin, D. H. P., Ken K-C Yeung, David F. Cechetto, Shawn N. Whitehead, Sublimation of DAN Matrix for the Detection and Visualization of Gangliosides in Rat Brain Tissue for MALDI Imaging Mass Spectrometry. J Vis Exp 2017, (121).

28. Hines, K. M.; May, J. C.; McLean, J. A.; Xu, L., Evaluation of Collision Cross Section Calibrants for Structural Analysis of Lipids by Traveling Wave Ion Mobility-Mass Spectrometry. Anal. Chem. 2016, 88 (14), 7329–36.

29. Ross, D. H.; Seguin, R. P.; Xu, L., Characterization of the Impact of Drug Metabolism on the Gas-Phase Structures of Drugs Using Ion Mobility-Mass Spectrometry. Anal. Chem. 2019, 91 (22), 14498–14507.

30. Ross, D. H.; Cho, J. H.; Xu, L., Breaking Down Structural Diversity for Comprehensive Prediction of Ion-Neutral Collision Cross Sections. Anal. Chem. 2020, 92 (6), 4548–4557.

31. Zhou, Z.; Luo, M.; Chen, X.; Yin, Y.; Xiong, X.; Wang, R.; Zhu, Z. J., Ion mobility collision cross-section atlas for known and unknown metabolite annotation in untargeted metabolomics. Nat Commun 2020, 11 (1), 4334.

32. Gregory W. Vandergrift, J. K. L., Michael J. Taylor, Kevin J. Zemaitis, Theodore Alexandrov, Josie G. Eder, Heather M. Olson, Jennifer E. Kyle, Christopher Anderton, Are Phosphatidic Acids Ubiquitous in Mammalian Tissues or Overemphasized in Mass Spectrometry Imaging Applications? Analysis & Sensing 2023.

33. Lee, R. W.; Conley, S. K.; Gropman, A.; Porter, F. D.; Baker, E. H., Brain magnetic resonance imaging findings in Smith-Lemli-Opitz syndrome. Am J Med Genet A 2013, 161A (10), 2407–19.

34. Picache, J. A.; Rose, B. S.; Balinski, A.; Leaptrot, K. L.; Sherrod, S. D.; May, J. C.; McLean, J. A., Collision cross section compendium to annotate and predict multi-omic compound identities. Chem Sci 2019, 10 (4), 983–993.

35. Dang Do, A. N.; Baker, E. H.; Warren, K. E.; Bianconi, S. E.; Porter, F. D., Spontaneously regressing brain lesions in Smith-Lemli-Opitz syndrome. Am J Med Genet A 2018, 176 (2), 386–390.

36. Le Belle, J. E.; Orozco, N. M.; Paucar, A. A.; Saxe, J. P.; Mottahedeh, J.; Pyle, A. D.; Wu, H.; Kornblum, H. I., Proliferative neural stem cells have high endogenous ROS levels that regulate self-renewal and neurogenesis in a PI3K/Akt-dependant manner. Cell Stem Cell 2011, 8 (1), 59–71.

37. Tan, S. T.; Ramesh, T.; Toh, X. R.; Nguyen, L. N., Emerging roles of lysophospholipids in health and disease. Prog Lipid Res 2020, 80, 101068.

38. Joensuu, M.; Wallis, T. P.; Saber, S. H.; Meunier, F. A., Phospholipases in neuronal function: A role in learning and memory? J. Neurochem. 2020, 153 (3), 300–333.

