## Supplemental Figures S1-S5 for "MALDI-IM-MS Imaging of Brain Sterols and Lipids in a Mouse Model of Smith-Lemli-Opitz Syndrome"

**Table of Content**

Table S1. Table of lipid identifications and measured CCS values compared to reference values (% error).

Table S2. Table of isobaric lipid pairs with different CCS values.

Figure S1. PCA and OPLS-DA plots.

Figure S2. Sterol and oxysterol images (biological replicate data for Figure 2).

Figure S3. Other isobaric lipid pairs with different CCS values.

Figure S4. Significantly altered lipid images (biological replicate data for Figure 5).

Figure S5. Other lipid species from S-plot.

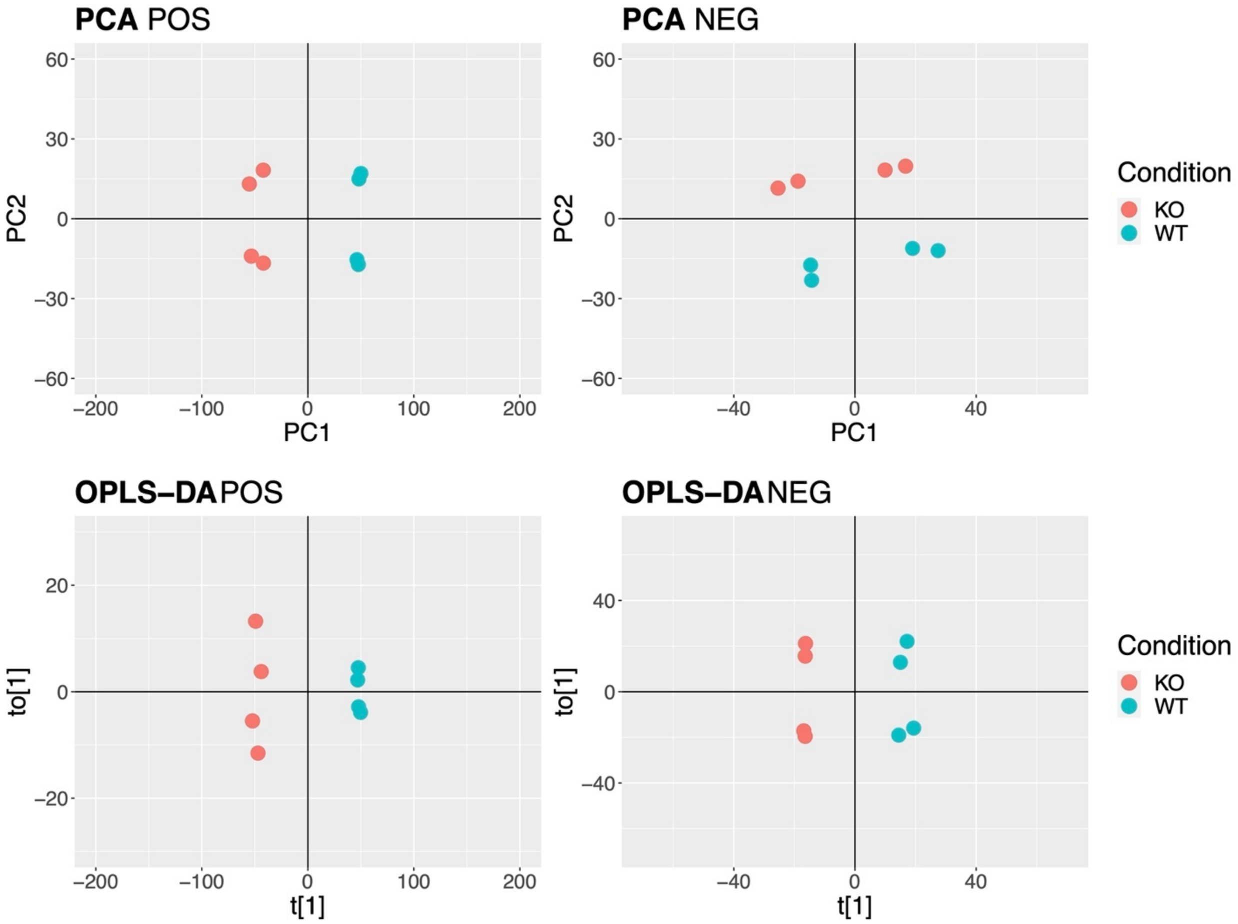


Figure S1. Multivariate statistical analysis of WT and *Dhcr7*-KO mouse brain tissues. Unsupervised PCA plots for one biological replicate for both negative and positive ionization modes. Supervised OPLS-DA plots.

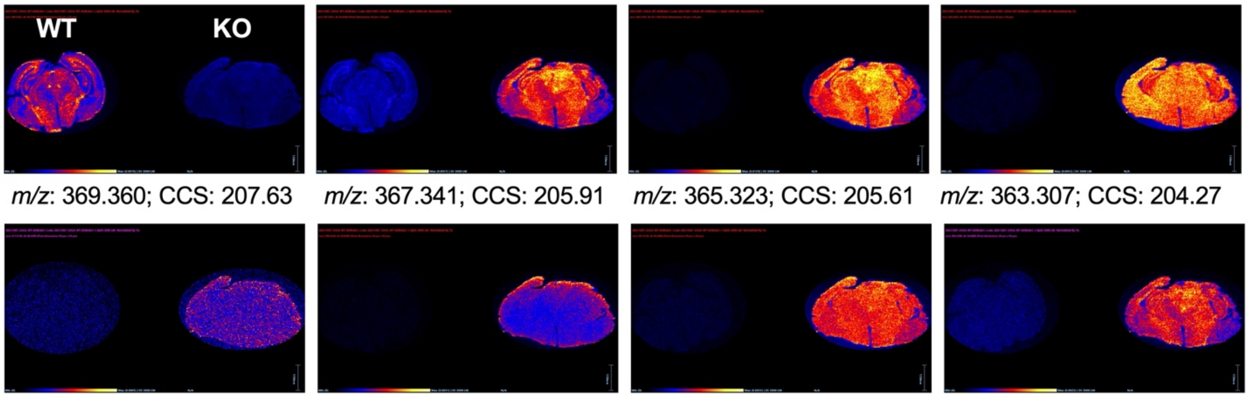


*m/z*: 369.360; CCS: 207.63 *m/z*: 367.341; CCS: 205.91 *m/z*: 365.323; CCS: 205.61 *m/z*: 363.307; CCS: 204.27

*m/z*: 417.325; CCS: 212.56 *m/z*: 399.329; CCS: 211.26 *m/z*: 381.318; CCS: 207.54 *m/z*: 383.334; CCS: 207.79

Figure S2. Biological replicate data for sterol and oxysterol features from Figure 2 (n = 2 animals per genotype and per ionization mode). Left: WT; right: *Dhcr7*-KO.

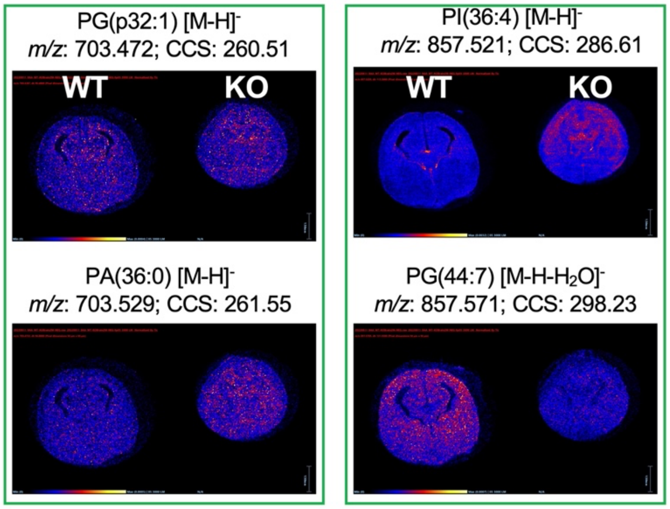


Figure S3. Other isobaric lipid pairs with different CCS values. Left: WT; right: *Dhcr7*-KO.

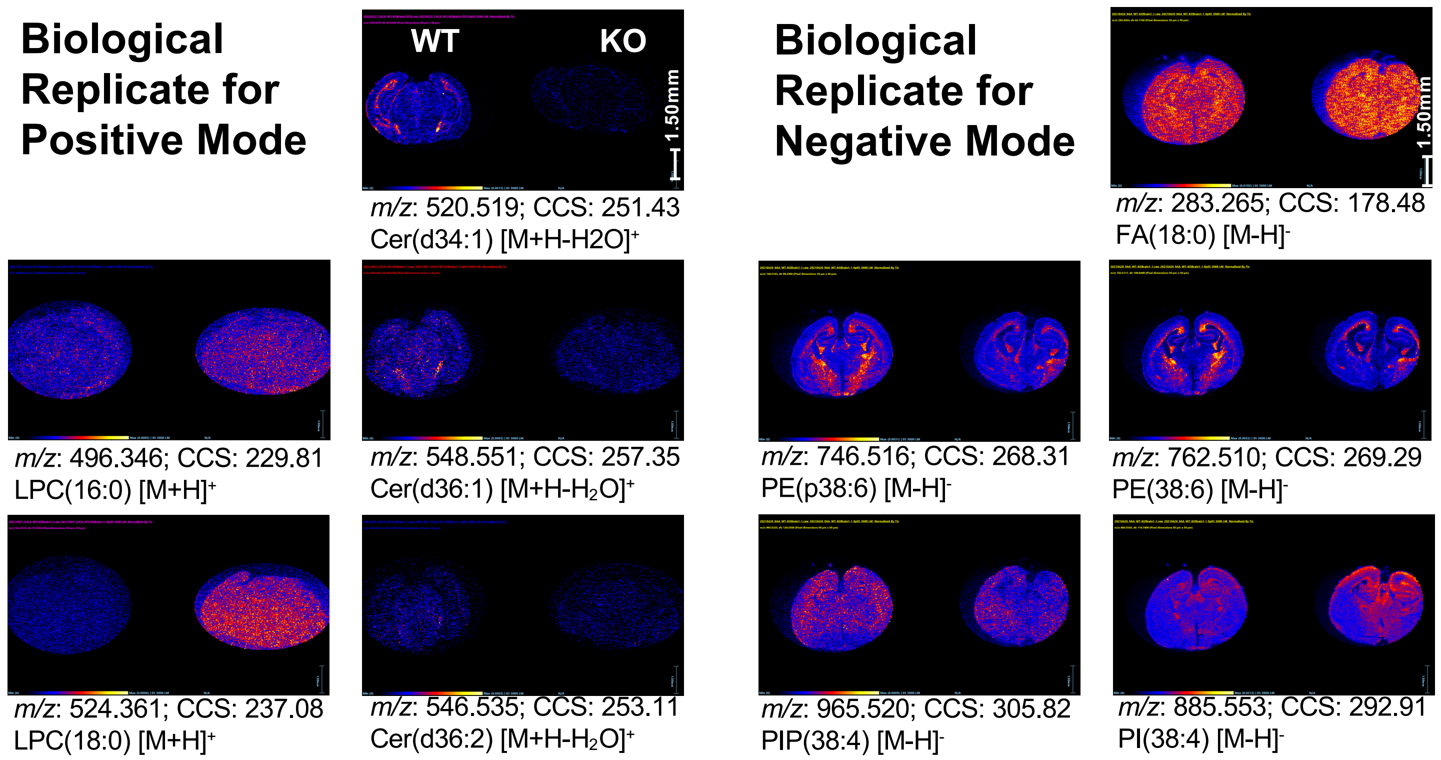


Figure S4. Biological replicate data for significantly altered lipid features from Figure 5 (n = 2 animals per genotype and per ionization mode). Left: WT; right: *Dhcr7*-KO.

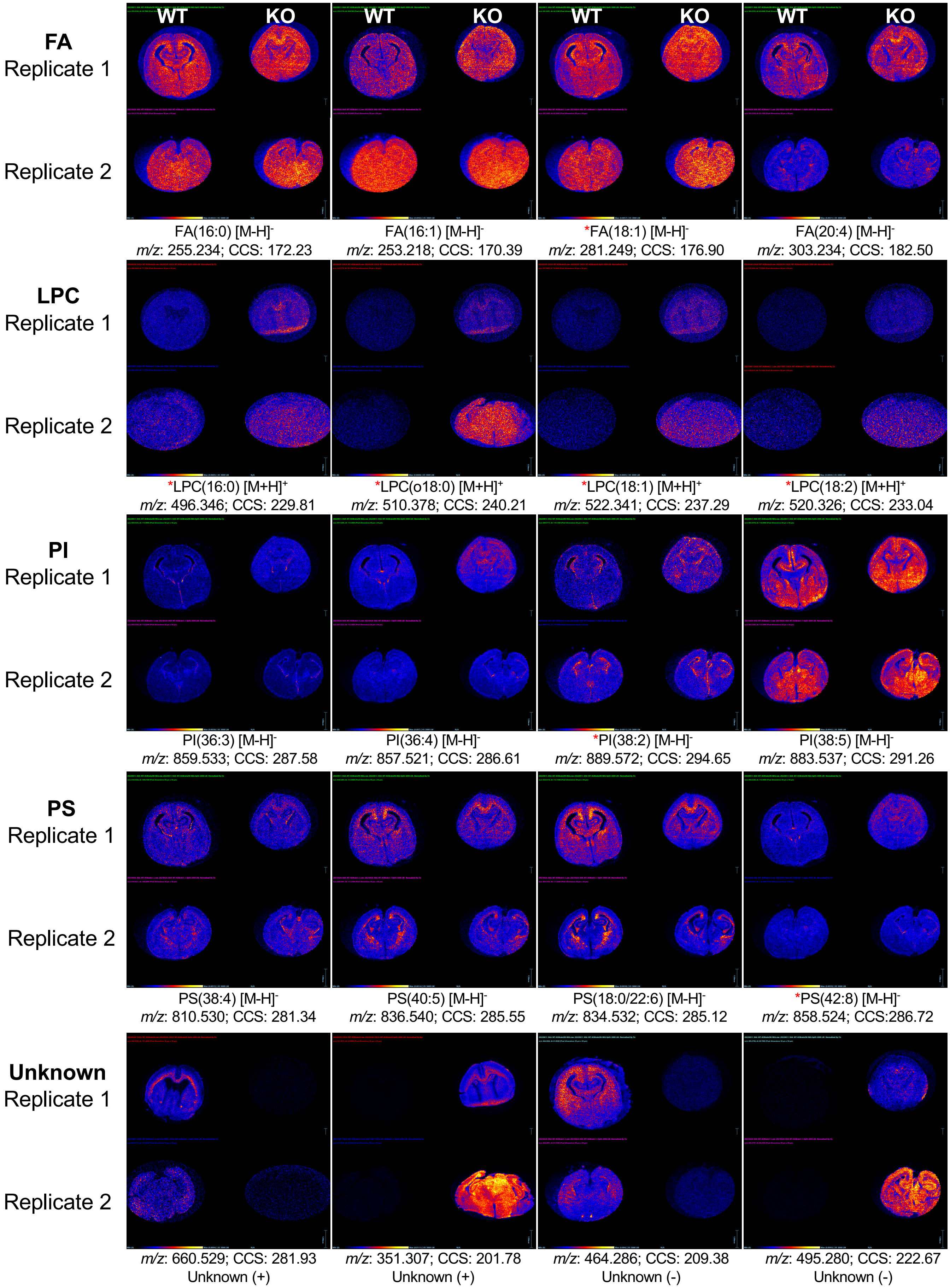


Figure S5. Additional lipid species that were significantly altered between WT (left) and *Dhcr7*-KO (right) brain sections (determined via S-plot analysis), including unidentified features.
